# Hypoxia on vhl-deficient cells to obtain hif related genes through bioinformatics analysis

**DOI:** 10.1101/863662

**Authors:** Hua Lin, Hongwei Liu

## Abstract

Vhl is responsible for degrading the transcription factor hif-1. Hif-1 transcription factors drive changes in hypoxic gene expression to adapt cells to exposure to hypoxic environments. This study hopes to analyze the effect of hypoxia on the vhl-deficient cells to obtain hif regulated genes through bioinformatics analysis. The top ten genes evaluated by connectivity degree in the PPI network were identified. CDK1, CTNNB1, NHP2, CCNA2, CTNNB1, MRPL16, CCND1 were down-regulated. CDK1, RPL12, RPL17, RPL27, RPS10 were up-regulated.

## Introduction

Vhl is responsible for degrading the transcription factor hif-1. Hif-1 transcription factors drive changes in hypoxic gene expression to adapt cells to exposure to hypoxic environments. Hypoxia active the heterodimeric transcription factor hif-1 to trigger a synergistic transcriptional response resulting in solid tumours.^3^ under oxygen deprivation hif1 is an essential helix-loop-helix PAS domain transcription factor.^4-7^ Hif-1α dimerizes with hif-1β in the nucleus, hif-1 decreased the speed of degrading under hypoxia condition. Hypoxia response elements are the heterodimer of hif-1 binding to regulatory DNA sequences. Transcriptional co-activators Promote angiogenesis through enhancing the transcription of a lot of target genes.^4,5,8^ Vhl gene is a tumour suppressor gene, and its proteins pvHL30 and pVHLl9. It has the function of tumour inhibition, and the two structures are similar, which are collectively called vhl protein(pvHL). Hypoxia-inducible factor hif gene is a target gene of vhl gene. hif is an oxygen-dependent transcriptional activator produced by cells during hypoxia, by only B subunits. Oxygen regulates hif activity primarily through hif-a. For cell growth, inhibition of hif1 expression can be promoted, causing the death of the cells under hypoxia condition. Therefore, this study hopes to analyze the effect of hypoxia on the vhl-deficient cells to obtain hif regulated genes through bioinformatics analysis.

## Materials and methods

We download a database from the GEO database(https://www.ncbi.nlm.nih.gov/geo/) to obtain the gene expression datasets. One series(GDS1772) was selected out from the database about human, three groups including control, hypoxia and hypoxia-reoxygenation were retrieved from the database. All of those three groups contain two parts of vhl minus and vhl plus.(Table 1) GDS1772 was based on the Agilent GPL3423: Stanford Denko EOS Human 35K GeneChip v1.1. All of the data we’re freely available online. This study has not been reported by any experiment on humans and declared by any other authors.

**Table 1.** Dividing into three groups according to the degree of hypoxia.

|  | First group | Second group | Third group |
| --- | --- | --- | --- |
| VHL minus | Control 1 | Hypoxia 1 | Hypoxia (1h or 4h) and reoxygenation 1 |
| VHL plus | Control 2 | Hypoxia 2 | Hypoxia (1h or 4h) and reoxygenation 2 |

### Data processing of DEGs

R: The R Project for Statistical Computing(https://www.r-project.org/) was used to detect the DEGs of three groups between vhl minus and vhl plus samples, and the adjusted P-value and |logFC| were calculated. We selected the DEGs by adjusting P<0.01 and |logFC|≥2.0.

### GO and KEGG pathway analysis of DEGs

GO analysis is a widely used method for functional enrichment studied. Also, gene functions were composed of biological process (BP), molecular function (MF), and cellular component (CC) three parts. KEGG is a large-scale used database, including vast amounts of genomes, biological pathways, diseases, chemicals, and drugs. We use the Database for Annotation, Visualization, and Integrated Discovery (DAVID) tools (https://david.ncifcrf.gov/) to analysis the DEGs through GO annotation analysis and KEGG pathway enrichment analysis. P<0.05 was considered statistically significant.^18^

### PPI network construction and hub gene identification

In this study, we use the Search Tool for the Retrieval of Interacting Genes (STRING) database (http://string-db.org/) and GeneMANIA online database (https://genemania.org/) to analyze the PPI information and evaluate the potential PPI relationship. They were also used to identify the DEGs to analysis the PPI information. A combined score was set to 0.4, and then the PPI network was visualized by Cytoscape software (www.cytoscape.org/). The stability of the entire system was guaranteed by a higher degree node of connectivity. We calculated the degree of each protein node by using CytoHubba, a plugin in Cytoscape. Through those steps, we can select ten hub genes.^18^

## Results

### Identification of DEGs

Gene expression profile (GDS1772) was selected in this study. GDS1772 contained vhl minus and vhl plus samples. Based on the criteria of P< 0.05 and |logFC|≥1, the first group of control without hypoxia and the third group hypoxia-reoxygenation have no differential genes with each other. The second group of hypoxia, we found that a total of 376 DEGs were identified from vhl minus samples compared with vhl plus samples, including 162 upregulated genes and 214 downregulated genes(Table 2).

**Table 2.**
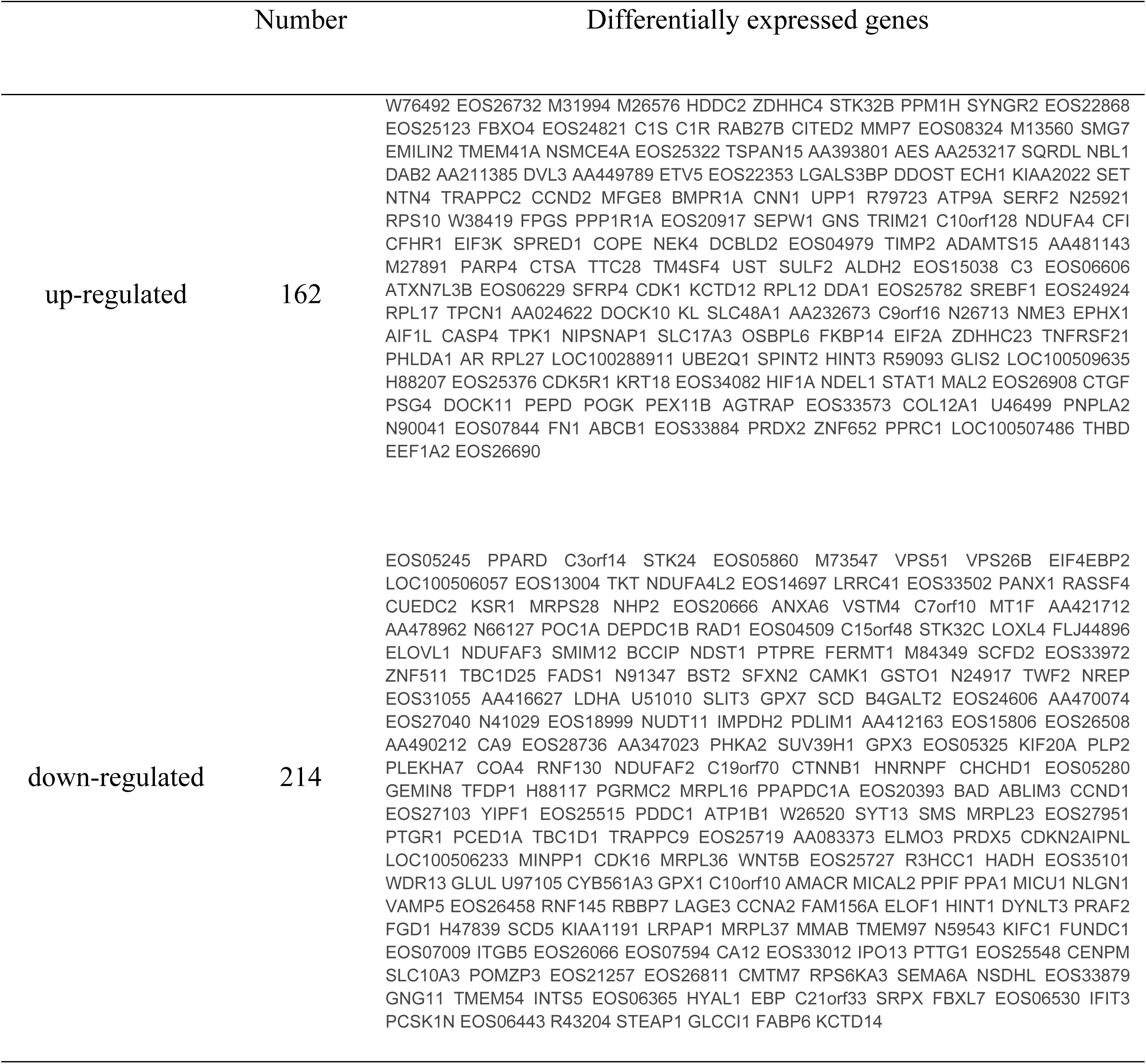
Differentially expressed genes.

### Functional enrichment analyses of DEGs

GO function and KEGG pathway enrichment analysis for DEGs were performed using the Cytoscape software(Table 3). The enriched GO terms were divided into CC, BP, and MF ontologies. BPs including cartilage development, oxidoreductase activity, acting on paired donors, with the oxidation of a pair of donors resulting in the reduction of molecular oxygen to two molecules of water and acyl-CoA desaturase activity. MF including large ribosomal subunit rRNA binding. CC, including mitochondrial protein complex. In addition, the results of KEGG pathway analysis showed that DEGs were mainly enriched in pathways in biosynthesis of unsaturated fatty acids (Table 4).

**Table 3.** The enrichment analysis of differentially expressed genes between vhl minus and vhl plus.

| Category | Term | PValue | Associated Genes Found |
| --- | --- | --- | --- |
| GOTERM_BP_DIRECT | GO:0051216~ cartilage development | 0.02687* | ANXA6, BMPR1A, CCN2, **CCNA2**, COL12A1, **CTNNB1**, HIF1A, HYAL1, POC1A, PRDX2, SULF2, WNT5B |
| GOTERM_BP_DIRECT | GO:0016717~ oxidoreductase activity, acting on paired donors, with oxidation of a pair of donors resulting in the reduction of molecular oxygen to two molecules of water | 0.00364* | FADS1, SCD, SCD5 |
| GOTERM_BP_DIRECT | GO:0016215~ acyl-CoA desaturase activity | 0.00148* | FADS1, SCD, SCD5 |
| GOTERM_CC_DIRECT | GO:0098798~ mitochondrial protein complex | 0.02403* | C15orf48, CHCHD1, HADH, MICOS13, MICU1, **MRPL16**, MRPL23, MRPL36, MRPL37, MRPS28, NDUFA4, NDUFA4L2, PPIF |
| GOTERM_MF_DIRECT | GO:0070180~ large ribosomal subunit rRNA binding | 0.01969* | MRPL23, **RPL12**, **RPL17** |

\*Pvalue<0.05

**Table 4.** The KEGG pathway enrichment analysis of DEG differentially expressed genes between vhl minus and vhl plus.

| Term | Pvalue | Associated Genes |
| --- | --- | --- |

|  |  | Found |
| --- | --- | --- |
| KEGG:01040~ Biosynthesis of unsaturated fatty acids | 0.03303* | ELOVL1, FADS1, SCD, SCD5 |

\*Pvalue<0.05

### PPI network construction and hub gene identification

Protein interactions among the DEGs were predicted with STRING tools and GeneMANIA online database. A total of 268 nodes and 428 edges were involved in the PPI network, as presented in Figure 1. The top ten genes evaluated by connectivity degree in the PPI network were identified. The results showed that ribosomal protein L12(RPL12) was the most outstanding gene with connectivity degree=12, followed by ribosomal protein L27(RPL27; degree=13), ribosomal protein L17(RPL17; degree=10), ribosomal protein S10(RPS10; degree=11), NHP2 ribonucleoprotein(NHP2; degree=10), cyclin D1(CCND1;degree=21), cyclin A2(CCNA2; degree=14), mitochondrial ribosomal protein L16(MRPL16; degree=10), cyclin-dependent kinase 1(CDK1; degree=15), catenin beta 1(CTNNB1; degree=20). All of these hub genes were upregulated in vhl plus.

**Figure 1.**
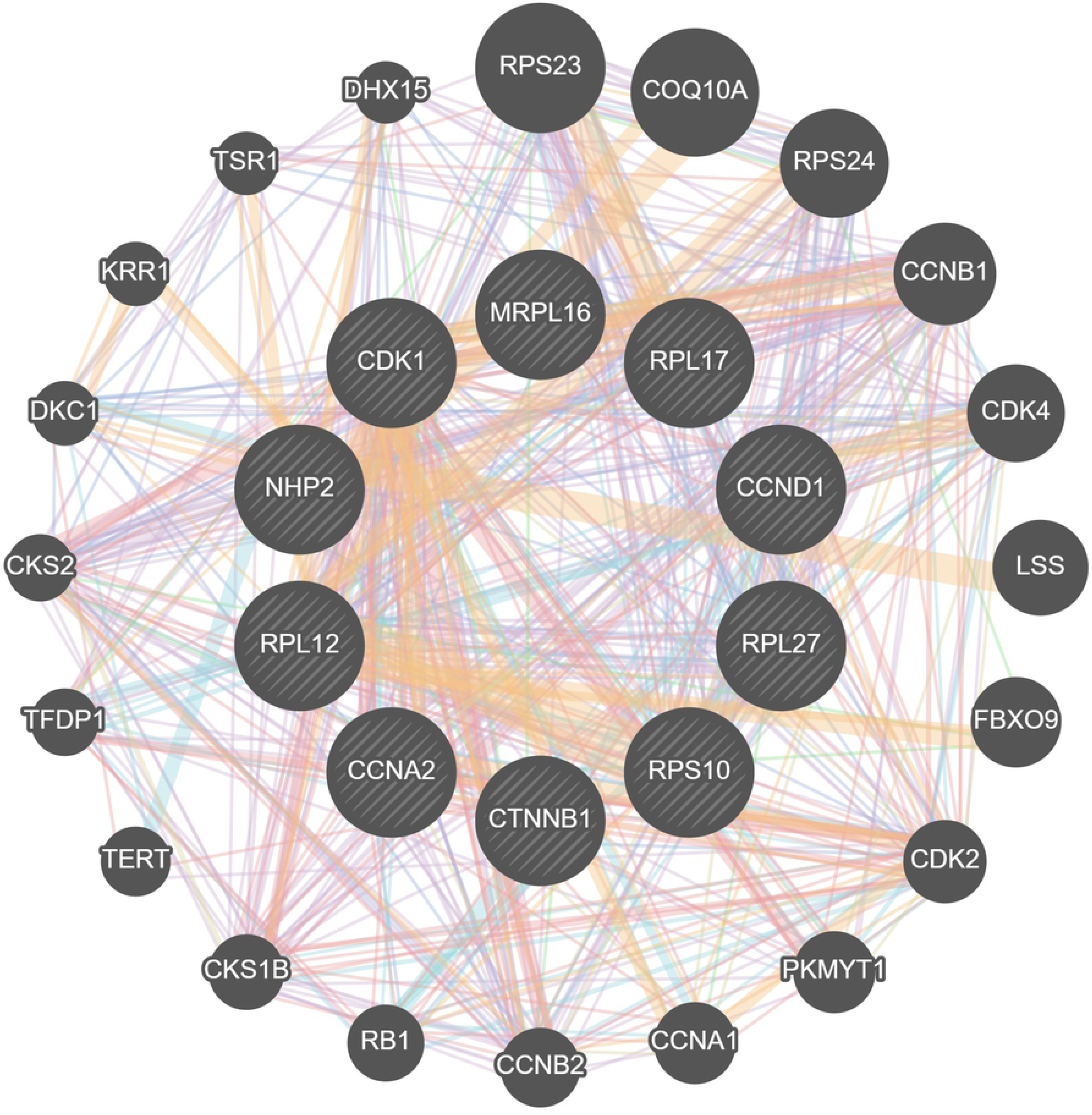
Based on STRING and GeneMANIA online database, protein-protein interaction networks of the differentially expressed genes were constructed and modular analyses

**Figure 2.**
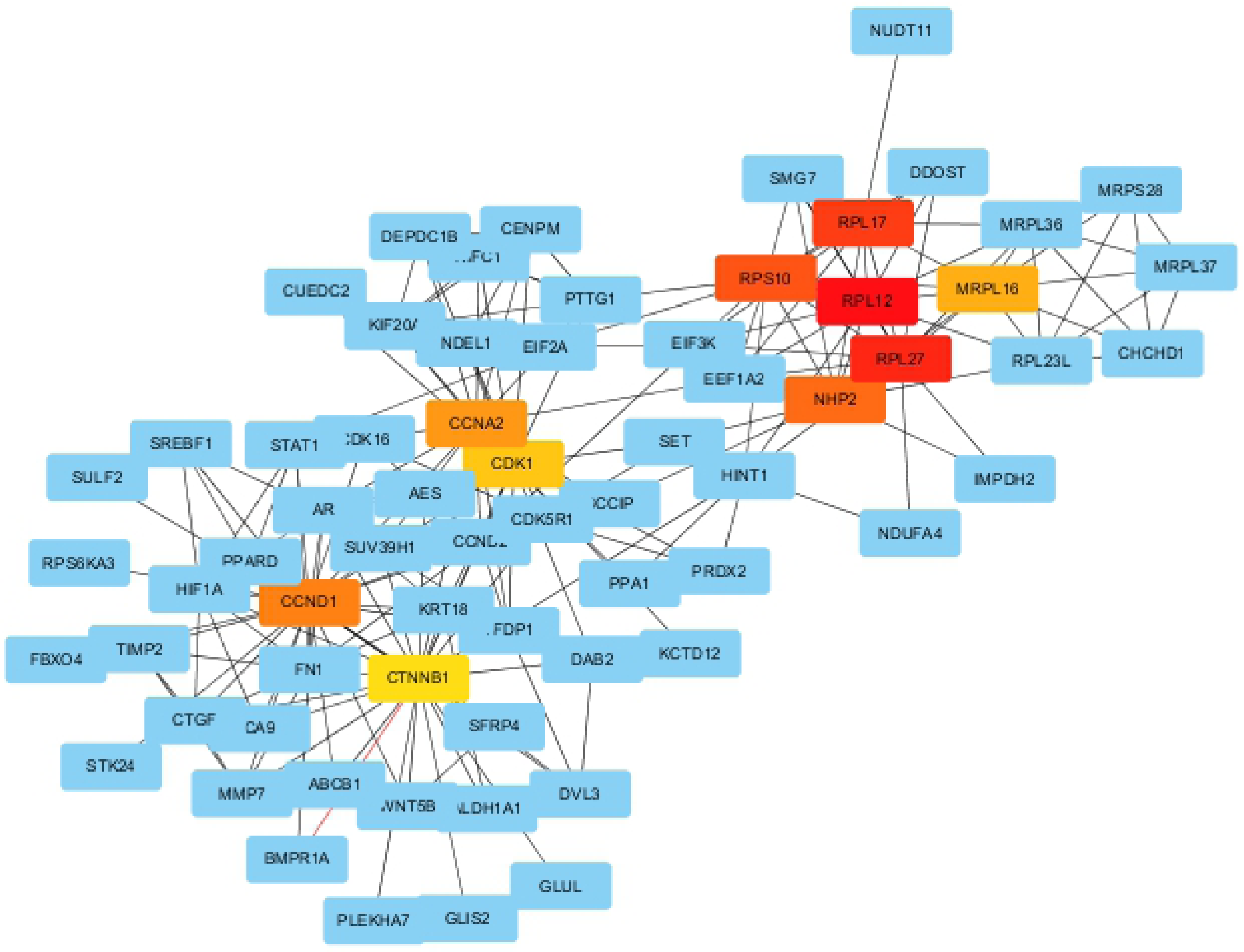
The PPI network was visualized by Cytoscape software and the top ten genes was evaluated by connectivity degree in the PPI network

## Discussion

In recent years, with the rapid development of modern biotechnology, such as biochip and high-throughput sequencing, bioinformatics has attracted more and more attention. Through big data analysis, it is an important application of bioinformatics to explore relevant genes that play a leading role in the development of diseases and provide ideas for the study of the mechanism of diseases. This study hopes to analyze the effect of hypoxia on vhl-deficient cells to obtain hif regulated genes through bioinformatics analysis. The first group of control without hypoxia and the third group hypoxia reoxygenation have no differential genes with each other. The second group of hypoxia, we found that a total of 376 DEGs were identified from vhl minus samples compared with vhl plus samples, including 162 up-regulate genes and 214 downregulated genes. The enriched GO terms were divided into CC, BP, and MF ontologies. BPs including cartilage development, oxidoreductase activity, acting on paired donors, with oxidation of a pair of donors resulting in the reduction of molecular oxygen to two molecules of water and acyl-CoA desaturase activity. MF including large ribosomal subunit rRNA binding. CC, including mitochondrial protein complex. In addition, the results of KEGG pathway analysis showed that DEGs were mainly enriched in pathways in biosynthesis of unsaturated fatty acids. The top ten genes evaluated by connectivity degree in the PPI network were RPL12, RPL27, RPL17, RPS10, NHP2, CCND1, CCNA2, MRPL16, CDK1, CTNNB1. NHP2, CCNA2, CTNNB1, MRPL16, CCND1 were down-regulated. CDK1, RPL12, RPL17, RPL27, RPS10 were up-regulate. According to the enrichment analysis of DEG between vhl minus and vhl plus, CCNA2 and CTNNB1 were related to cartilage development. MRPL16 had corresponded with a mitochondrial protein complex. RPL12 and RPL17 involved in large ribosomal subunit rRNA binding.

CCNA2, also called cyclin A2. The protein encoded by CCNA2 is part of the Highly conservative cyclin family. Cyclin family members operate as regulating the cell cycle, which can bind and active cyclin-dependent kinase2 to promote the transition through G1/S and G2/M. Deng H, et al. find that overexpressed CyclinA2 improve the cardiomyocytes proliferation under hypoxia impair. Other articles find that in the neonatal rat hearts, the proliferation of cardiomyocytes was inhibited under hypoxia. However, overexpressed CyclinA2 weaken the proliferation of cardiomyocytes suggesting that CyclinA2 is an essential part of regulating cardiomyocytes growth and increasing protection of cardiomyocytes from disability of proliferation in hypoxic conditions.^9^ A few days after birth, the proliferative function of mammals’ cardiomyocytes almost lost because of the sharp down-regulation of CyclinA2 in the heart.^10^ Previous studies indicated that in the adult heart CyclinA2 almost have no function. Increasing the content of CyclinA2 in adult animals’ heart can make a difference in restoration of cardiac proliferation.^10,11^ Also, Schulze A, et al. find that in the primary neonatal rat cardiomyocytes overexpressed CyclinA2 increase proliferation. After exposure to hypoxia for 12h, CyclinA2 down-regulate and marked decrease the cardiomyocyte proliferation. In conclusion, overexpressed CyclinA2 have beneficial to cardiomyocyte growth under impair of hypoxia.^12^ In summary, decrease the expression of vhl can up-regulate CyclinA2 and also increase hif. There was no other organs protection related studied about CyclinA2 protection under hypoxia.

CDK1, also called cyclin dependent kinase 1. The protein encoded by CDK1 is belonging to the Ser/Thr protein kinase family. It is a catalytic subunit of M-phase promoting factor(MPF), and have an essential function in the transition of the eukaryotic cell cycle for G1/S and G2/M phase. This protein is related to mitotic cyclins stably, which has a function to regulate subunits. The content of cyclin and the phosphorylation and dephosphorylation of this protein in the cell cycle can regulate the kinase activity of this protein. This gene has spliced transcript variants which encodes various isoforms. The protein encoded by CDK1 belongs to the Ser/Thr protein kinase family and also is a catalytic submit of M-phase promoting factor(MPF). MPF is important for the transition of G1/S and G2/M in the eukaryote cell cycle. This protein is regulatory subunit related to Mirotic cycling stable. Therefore, no matter whether hypoxia, CDK1 regulate the phosphorylation of hif-1a, which works for the steady-state levels of hif1a. Inhibitors of CDK1 have been suggested using for the treatment of many kinds of cancers and that it can increase the sensitization of the TRAIL-induced apoptosis.^15-17^ Inhibiting CDK1 can decrease hif-1a expression, and active transcription and decrease surviving phosphorylation so that increasing the sensitivity of cancer cells apoptosis. Recently, Herrera MC, et al find that regulating RNA polymerase III activity affects cdk1 gates cell cycle-dependent tRNA synthesis.^19^ Jones MC, et al. find cdk1 also related to cell adhesion.^20^ Wang Z, et al. find that during the cell cycle G2/M Progression, Cdk1 coordinates Mitochondrial Respiration.^22^

So that, vhl can increase the expression of CDK1, and CDK1can regulate and activate the expression of hif-1a under the condition of hypoxia. CDK1 was mainly studied in the regulation of cancer cells apoptosis at present.

CTNNB1, also called catenin beta 1. The protein encoded by CTNNB1 belongs to the constitute adherents junctions (AJs), which can regulate cell growth and adhere between cells to create and maintain epithelial cell layers. This protein works on acting cytoskeleton and transmitting the contact inhibition signal to stop dividing once the epithelium is formed. A kind of cancer can be caused by a mutation of this gene. We suppose that this gene inhibits the dividing of cancer cells.^1,24^ Hirata H, et al. find that β-catenin promotes cell survival and adapt to hypoxia by strengthening hif-1 mediated transcription. So that, we find that vhl down-regulate the expression of CTNNB1 same as the result of Hirata H’s findings and CTNNB1 can protect cells from hypoxia.^1^

NHP2, CCNA2, CTNNB1, MRPL16, CCND1 were down-regulated by vhl. NHP2 has an effect on the rRNA production and rRNA pseudouridylation. MRPL16 the protein encoded by MRPL16 comprises the mitoribosome. Mutations, amplification and overexpression of CCND1 alters cell cycle progression, which is observed in various tumors and conduces to tumorigenesis. Recent article find that targeting USP22 and CDK inhibitors have a benefit on cancer patients induced by CCND1.21 Xu P, et al. find thatCCND1 has the disadvantage of early-stage lung adenocarcinoma patients.23 CDK1, RPL12, RPL17, RPL27, RPS10 were up-regulated by vhl. The expression of RPS10 was related to colorectal cancers. RPL12, RPL17 and RPL27 are related to encoding the ribosomal protein. In the condition of hypoxia, other genes affecting the expression of hif have not been reported, which would be the direction of our follow-up study.

## Acknowledgments

The authors would like to thank Jiajun Han for his supporting.

## Reference

1. Hirata H, Hinoda Y, Ueno K, Nakajima K, Ishii N, Dahiya R. MicroRNA-1826 directly targets beta-catenin (CTNNB1) and MEK1 (MAP2K1) in VHL-inactivatedrenal cancer. Carcinogenesis. 2012; 33(3):501–8.doi: 10.1093/carcin/bgr302

2. Pirrotta MT, Bernardeschi P, Fiorentini G. Targeted-therapy in advanced renal cell carcinoma. Curr Med Chem. 2011;18(11):1651–7.

3. Trang P, Weidhaas JB, Slack FJ. MicroRNAs as potential cancer therapeutics.. Oncogene. 2008; 27(12):S52–7. doi: 10.1038/onc.2009.353.

4. Inui M, Martello G, Piccolo S. MicroRNA control of signal transduction. 2010; 11(4):252–63. doi: 10.1038/nrm2868

5. Kohno M, Pouyssegur J. Targeting the ERK signaling pathway in cancer therapy. Ann Med. 2006;38(3):200–11.

6. Wang GL, Jiang BH, Rue EA, Semenza GL. Hypoxia-inducible factor 1 is a basic-helix-loop-helix-PAS heterodimer regulated by cellular O2tension. Proc Natl Acad Sci U S A. 1995; 92(12):5510–4.

7. Wenger RH, Stiehl DP, Camenisch G. Integration of oxygen signaling at the consensus HRE. Sci STKE. 2005;18(10):re12.

8. Rankin EB, Giaccia AJ. The role of hypoxia-inducible factors in tumorigenesis. Cell Death Differ 2008; 15:678-85. doi. org/10.1038/cdd.2008.21

9. Deng H, Cheng Y, Guo Z, Zhang F, Lu X, Feng L, et al. Overexpression of CyclinA2 ameliorates hypoxia-impaired proliferation of cardiomyocytes. Exp Ther Med. 2014;8(5):1513–1517.

10. Chaudhry HW, Dashoush NH, Tang H, Zhang L, Wang X, Wu EX, et al. Cyclin A2 mediates cardiomyocyte mitosis in the postmitotic myocardium. J Biol Chem. 2004;20(8):35858–66.

11. Woo YJ, Panlilio CM, Cheng RK, Liao GP, Atluri P, Hsu VM, et al. Therapeutic delivery of cyclin A2 induces myocardial regeneration and enhances cardiacf unction in ischemic heart failure. Circulation. 2006;4(7):I206–13.

12. Schulze A, Zerfass K, Spitkovsky D, Middendorp S, Bergès J, Helin K, et al. Cell cycle regulation of the cyclin A gene promoter is mediated by a variant E2F site. Proc Natl Acad Sci U S A. 1995;21(11):11264–8.

13. Mayes PA, Dolloff NG, Daniel CJ, Liu JJ, Hart LS, Kuribayashi K, et al. Overcoming hypoxia-induced apoptotic resistance through combinatorial inhibition of GSK-3β and CDK1. Cancer Res. 2011;1(8):5265–75. doi: 10.1158/0008-5472.CAN-11-1383

14. Shapiro GI. Cyclin-dependent kinase pathways as targets for cancer treatment. J Clin Oncol. 2006;10(4):1770–83.

15. Kim DM, Koo SY, Jeon K, Kim MH, Lee J, Hong CY, Jeong S. Rapid induction of apoptosis by combination of flavopiridol and tumor necrosis factor (TNF)-alpha or TNF-related apoptosis-inducing ligand in human cancer cell lines. CancerRes. 2003;63(3):621–6.

16. Kim EH, Kim SU, Shin DY, Choi KS. Roscovitine sensitizes glioma cells to TRAIL-mediated apoptosis by downregulation of s urvivinand XIAP. Oncogene. 2004; 23(2):446–56.

17. Goga A, Yang D, Tward AD, Morgan DO, Bishop JM. Inhibition of CDK1 as a potential therapy for tumors over-expressing MYC. Nat Med. 2007;13(7):820–7.

18. Bu X, Wang B, Wang Y, Wang Z, Gong C, Qi F, et al. Pathway-related modules involved in the application of sevoflurane or propofol in off-pump coronary artery bypass graft. Exp Ther Med. 2017;14:97–106. doi: 10.3892/etm.2017.4504.

19. Herrera MC, Chymkowitch P, Robertson JM, Eriksson J, Bøe SO, Alseth I, et al. Cdk1 gates cell cycle-dependent tRNA synthesis by regulating RNA polymerase III activity. Nucleic Acids Res. 2018; 46(22):11698–11711. doi: 10.1093/nar/gky846

20. Jones MC, Askari JA, Humphries JD, Humphries MJ. Cell adhesion is regulated by CDK1 during the cell cycle. JCellBiol. 2018;217(9):3203–3218. doi: 10.1083/jcb.201802088

21. Gennaro VJ, Stanek TJ, Peck AR, Sun Y, Wang F, Qie S, et al. Control of CCND1 ubiquitylation by the catalytic SAGA subunit USP22 is essential for cell cycleprogression through G1 in cancer cells. Proc Natl Acad Sci U S A. 2018;115(40):E9298–E9307. doi: 10.1073/pnas.1807704115

22. Wang Z, Fan M, Candas D, Zhang TQ, Qin L, Eldridge A, et al. Cyclin B1/Cdk1 coordinates mitochondrial respiration for cell-cycle G2/M progression. Dev Cell. 2014; 29(2):217–32. doi: 10.1016/j.devcel.2014.03.012

23. Xu P, Zhao M, Liu Z, Liu Y, Chen Y, Luo R, et al. Elevated nuclear CCND1 expression confers an unfavorable prognosis for early stage lung adenocarcinoma patients. Int J Clin Exp Pathol. 2015; 8(12):15887–94

24. Wen J, Min X, Shen M, Hua Q, Han Y, Zhao L, et al. ACLY facilitates colon cancer cell metastasis by CTNNB1. J Exp Clin Cancer Res. 2019;38(1):401. doi: 10.1186/s13046-019-1391-9.

